# A novel single-domain Na^+^-selective voltage-gated channel in photosynthetic eukaryotes

**DOI:** 10.1101/2020.04.29.068528

**Authors:** Katherine E. Helliwell, Abdul Chrachri, Julie Koester, Susan Wharam, Alison R. Taylor, Glen L. Wheeler, Colin Brownlee

## Abstract

The evolution of Na^+^-selective four-domain voltage-gated channels (4D-Na_v_s) in animals allowed rapid Na^+^-dependent electrical excitability, and enabled the development of sophisticated systems for rapid and long-range signalling. Whilst bacteria encode single-domain Na^+^-selective voltage-gated channels (BacNa_v_), they typically exhibit much slower kinetics than 4D-Na_v_s, and are not thought to have crossed the prokaryote-eukaryote boundary. As such, the capacity for rapid Na^+^-selective signalling is considered to be confined to certain animal taxa, and absent from photosynthetic eukaryotes. Certainly, in land plants, such as the Venus Flytrap where fast electrical excitability has been described, this is most likely based on fast anion channels. Here, we report a unique class of eukaryotic Na^+^-selective single-domain channels (EukCatBs) that are present primarily in haptophyte algae, including the ecologically important calcifying coccolithophores. The EukCatB channels exhibit very rapid voltage-dependent activation and inactivation kinetics, and sensitivity to the highly selective 4D-Na_v_ blocker tetrodotoxin. The results demonstrate that the capacity for rapid Na^+^-based signalling in eukaryotes is not restricted to animals or to the presence of 4D-Na_v_s. The EukCatB channels therefore represent an independent evolution of fast Na^+^-based electrical signalling in eukaryotes that likely contribute to sophisticated cellular control mechanisms operating on very short time scales in unicellular algae.

**One Sentence Summary:** The capacity for rapid Na^+^-based signalling has evolved in ecologically important coccolithophore species via a novel class of voltage-gated Na^+^ channels, EukCatBs.

## Introduction

Electrical signals trigger rapid physiological events that underpin an array of fundamental processes in eukaryotes, from contractile amoeboid locomotion (Bingley and Thompson, 1962), to the action potentials of mammalian nerve and muscle cells (Hodgkin and Huxley, 1952). These events are mediated by voltage-gated ion channels (Brunet and Arendt, 2015). In excitable animal cells, Ca^2+^-or Na^+^-selective members of the four-domain voltage-gated cation channel family (4D-Ca_v_/Na_v_) underpin well-characterised signalling processes (Catterall et al., 2017). The 4D-Ca_v_/Na_v_ family is broadly distributed across eukaryotes, contributing to signalling processes associated with motility in some unicellular protist and microalgal species (Fujiu et al., 2009; Lodh et al., 2016), although these channels are absent from land plants (Edel et al., 2017). It is likely that the ancestral 4D-Ca_v_/Na_v_ channel was Ca^2+^ permeable, with Na^+^-selective channels arising later within the animal lineage (Moran et al., 2015). This shift in ion selectivity represented an important innovation in animals, allowing rapid voltage-driven electrical excitability to be decoupled from intracellular Ca^2+^ signalling processes (Moran et al., 2015).

Na^+^-selective voltage-gated channels have not been described in other eukaryotes, although a large family of Na^+^-selective channels (BacNa_v_) is present in prokaryotes (Ren et al., 2001; Koishi et al., 2004). BacNa_v_ are single-domain channels that form homotetramers, resembling the four-domain architecture of 4D-Ca_v_/Na_v_. Studies of BacNa_v_ channels have provided considerable insight into the mechanisms of gating and selectivity in voltage-dependent ion channels (Payandeh et al., 2012; Zhang et al., 2012). The range of activation and inactivation kinetics of native BacNa_v_ are generally slower than observed for 4D-Na_v_, suggesting that the concatenation and subsequent differentiation of individual pore-forming subunits may have enabled 4D-Na_v_ to develop specific properties such as fast inactivation, which is mediated by the conserved intracellular isoleucine–phenylalanine–methionine (IFM) linker between domains III and IV (Irie et al., 2010; Catterall et al., 2017) (**Figure 1A**).

**Figure 1.**
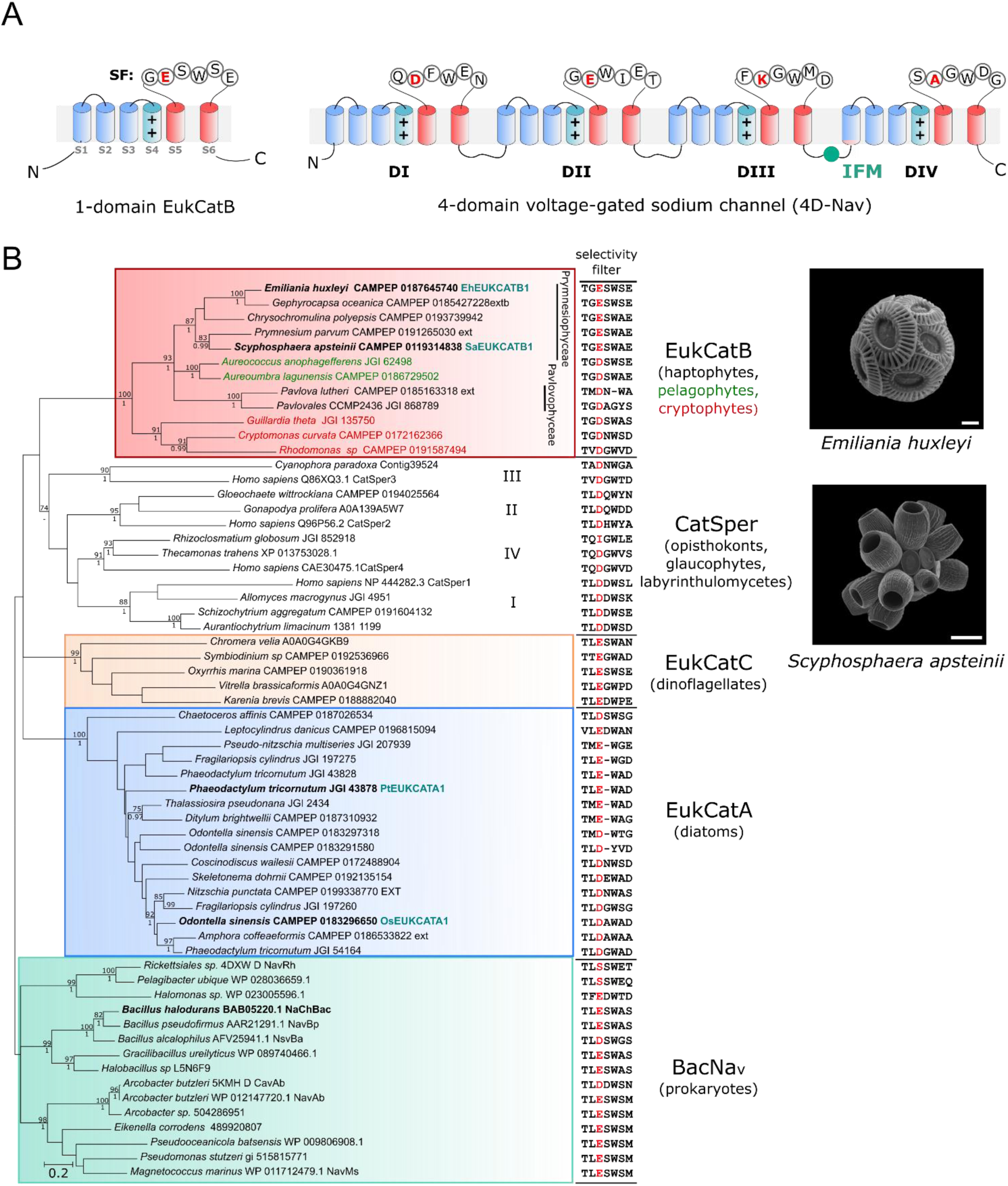
EukCatBs represent a novel class of single-domain channels. **A**. Schematic diagram of a single-domain EukCatB channel. The voltage-sensing module (S1-S4, blue) including conserved positively charged (++) residues of segment (S4) that responds to changes in membrane potential is shown. The pore module (S5-S6, red) is also indicated, including the selectivity filter (SF) motif (Ren et al., 2001). The structure of a 4D-Na_v_ (showing the selectivity filter of rat 4D-Na_v_1.4 with canonical ‘DEKA’ locus of Na^+^ selective 4D-Na_v_1s) is also displayed (right). The IFM motif of the fast inactivation gate is indicated (West et al., 1992) **B**. Maximum likelihood phylogenetic tree of single-domain voltage-gated channels including BacNa_v_ and the three distinct classes of EukCat channels (EukCatA-C). Representatives of the specialised family of single-domain Ca^2+^ channels identified in mammalian sperm (CatSpers) are also included. SF for each sequence is shown (right). “Position 0” of the high field strength site (HFS) that is known to be important in determining Na^+^ selectivity (Payandeh et al., 2011), is coloured red. Channel sequences selected for functional characterisation in this study are shown in bold. EukCatA sequences previously characterised in (Helliwell et al., 2019) are also indicated, as is NaChBac channel from *Bacillus halodurans* (Ren et al., 2001). ML bootstrap values (>70) and Bayesian posterior probabilities (>0.95) are above and below nodes, respectively. Scanning electron micrographs of coccolithophores *Emiliania huxleyi* (bar = 2 µm) and *Scyphosphaera apsteinii*, (bar = 10 µm) are shown.

We recently identified several classes of ion channel (EukCats) bearing similarity to BacNa_v_ in the genomes of eukaryotic phytoplankton. Characterisation of EukCatAs found in marine diatoms demonstrated that these voltage-gated channels are non-selective (exhibiting permeability to both Na^+^ and Ca^2+^) and play a role in depolarisation-activated Ca^2+^ signalling (Helliwell et al., 2019). Two other distinct classes of single-domain channels (EukCatBs and EukCatCs) were also identified that remain uncharacterised. These channels are present in haptophytes, pelagophytes and cryptophytes (EukCatBs), as well as dinoflagellates (EukCatCs) (Helliwell et al., 2019). Although there is a degree of sequence similarity between the distinct EukCat clades, the relationships between clades are not well resolved, and there is not clear support for a monophyletic origin of EukCats. The diverse classes of EukCats may therefore exhibit significant functional differences. Characterisation of these different classes of eukaryote single-domain channels is thus vital to our understanding of eukaryote ion channel structure, function and evolution, and to gain insight into eukaryote membrane physiology more broadly.

Notably, EukCatB channels were found in ecologically important coccolithophores, a group of unicellular haptophyte algae that represent major primary producers in marine ecosystems. Coccolithophores are characterised by their ability to produce a cell covering of ornate calcium carbonate platelets (coccoliths) (Taylor et al., 2017) (**Figure 1B**). The calcification process plays an important role in global carbon cycling, with the sinking of coccoliths representing a major flux of carbon to the deep ocean. Patch-clamp studies of coccolithophores indicate several unusual aspects of membrane physiology, such as an inwardly rectifying Cl^-^ conductance and a large outward H^+^ conductance at positive membrane potentials, which may relate to the increased requirement for pH homeostasis associated with intracellular calcification. Here we report that EukCatB channels from two coccolithophore species (*Emiliania huxleyi* and *Scyphosphaera apsteinii*) act as ultra-fast Na^+^-selective voltage-gated channels that exhibit many similarities to the 4D-Na_v_s that underpin neuronal signalling in animals. Thus, our findings demonstrate that the capacity for rapid Na^+^-based signalling has evolved in certain photosynthetic eukaryotes, contrary to previous widely held thinking.

## Results

### Coccolithophore EukCatBs are fast Na+-selective voltage-gated channels

EukCatB sequences are present in haptophyte, cryptophyte and pelagophyte taxa (Helliwell et al., 2019). They form a distinct phylogenetic group from the non-selective Ca^2+^ and Na^+^ permeable diatom EukCatAs and prokaryote BacNa_v_s (**Figure 1B; Supplementary Table 1**). Ion selectivity of voltage-gated channels is mediated by the conserved pore loop region between transmembrane segments S5 and S6, known as the selectivity filter (SF) (Catterall et al., 2017). Six amino acids (L<u>E</u>SWAS) make up this motif in the Na^+^-selective *Bacillus halodurans* BacNa_v_ channel (Yue et al., 2002) (species highlighted in bold, **Figure 1B**). Examination of the SFs of EukCat sequences indicates a clear diversification between the major EukCat clades. The SFs of coccolithophore (Prymnesiophyceae) EukCatB channels are strongly conserved (**Figure 1B**) and exhibit high similarity to Na^+^-selective BacNa_v_ channels. This includes the charged Glu (E) residue occupying “position 0” in the high field strength site (HFS) (highlighted red, **Figure 1B**) that is known to be important in determining Na^+^ selectivity (Payandeh et al., 2011). The adjacent Ser (S) and Trp (W) residues are also conserved. By comparison, EukCatAs from the diatoms, which we showed previously to be permeable to Na^+^ and Ca^2+^ (Helliwell et al., 2019), differ significantly from the canonical Na^+^ selective SF of NaChBac. EukCatAs either have an Asp (D) in position 0, or the conserved Glu (E) but not the adjacent Ser (S). Finally, dinoflagellate EukCatC sequences were highly diverse, ranging from SFs that were similar to the canonical Na^+^ selective motif (e.g. LESWSE, *Oxyrrhis marina* CAMPEP 0190361918) to SFs that resemble the Ca^2+^-selective NaChBac mutants (e.g. LEDWPE, *Karenia brevis* CAMPEP0188882040) (DeCaen et al., 2014).

To determine the ion transport properties of coccolithophore EukCatB channels, we expressed representative channels from *Emiliania huxleyi* (EhEUKCATB1) and *Scyphosphaera apsteinii* (SaEUKCATB1) in HEK293 cells (**Supplementary Table 2**). Patch clamp approaches revealed that both proteins yielded robust voltage-dependent inward currents that were distinct from those recorded with non-transfected cells (He and Soderlund, 2010) (**Supplementary Figure 1**). EhEUKCATB1 and SaEUKCATB1 were activated by membrane depolarisation, with extremely rapid activation and inactivation kinetics (**Figure 2; Supplementary Figure 2**). The kinetics are faster than those previously reported for native BacNa_v_s (Irie et al., 2010) (**Supplementary Table 3**) and are remarkably similar to those observed in animal 4D-Na_v_1s (τ_activation_ < 2 ms; τ_inactivation_ < 10 ms) (Hille, 2001; Ren et al., 2001).

**Figure 2.**
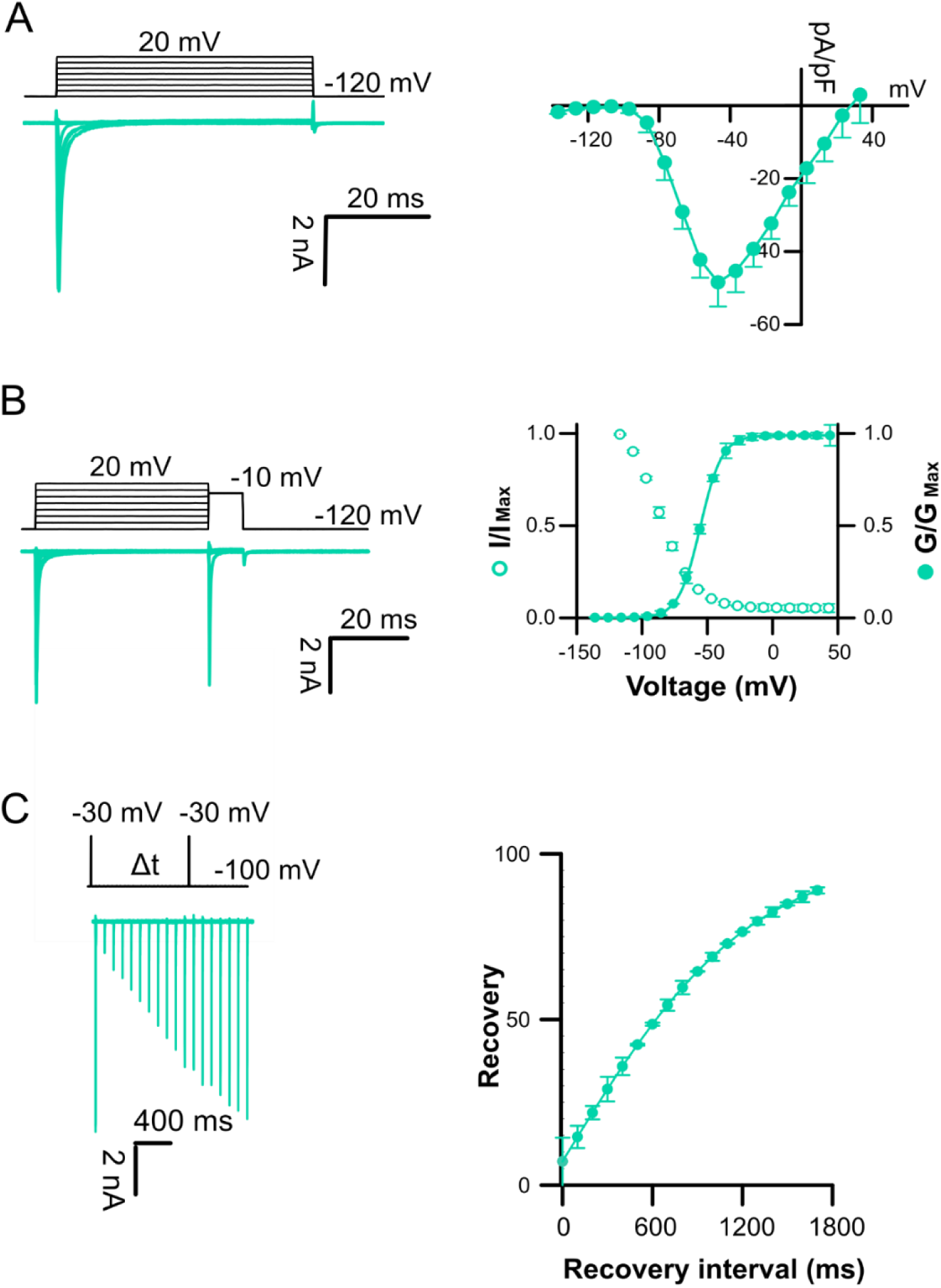
Coccolithophore EukCatB channels expressed in HEK293 cells yield currents with ultrafast kinetics. **A**. Typical current traces for cells expressing EhEUKCATB1 in response to membrane depolarisation. Average peak current-voltage (I-V) curves (right), where current was normalised to cell capacitance. Peak current = −57.6 ± 11.1 pA/pF (*n* = 10). **B**. Steady state inactivation: representative examples of current traces (left) used to obtain steady-state inactivation curves (right) for EhEUKCATB1 (*n* = 14). Average normalized data were fitted using the Boltzmann equation (see methods) for both activation (solid circles) and steady state inactivation (open circles) curves. Error bars: SEM (**see Supplementary Table 3**). **C**. Recovery from inactivation; superimposed currents obtained by a double-pulse protocol using a varying interval (Δt) between the two voltage pulses (left). Holding potential was −100 mV and test potential was −30 mV for 5 ms followed by a recovery test pulse to −30 mV for 5 ms between 100-1800 ms after the first test pulse. The peak currents elicited by the recovery pulse were normalised in order to construct the recovery curve. A single exponential was fitted to the averaged normalised recovery curve yielding τ for recovery (first order exponential fit) of 632 ± 50.6 (*n*=14).

To determine the ion selectivity of EukCatB channels, we systematically replaced cations in the external solution. Omitting Na^+^ from the external solution (substituting with 140 mM N-methyl-D-glutamine (NMDG), a large synthetic cation) completely inhibited the inward currents SaEUKCATB1, indicating that they have an absolute requirement for Na^+^ (**Figure 3A; Supplementary Table S4**). In contrast, the removal of Ca^2+^ from the external solution significanly enhanced the channel conductance in SaEUKCATB1. These data suggest that SaEUKCATB1 represents a fast-activating Na^+^-selective channel that is entirely distinct from other Na^+^-selective channels previously described in eukaryotes (4D-Na_v_).

**Figure 3.**
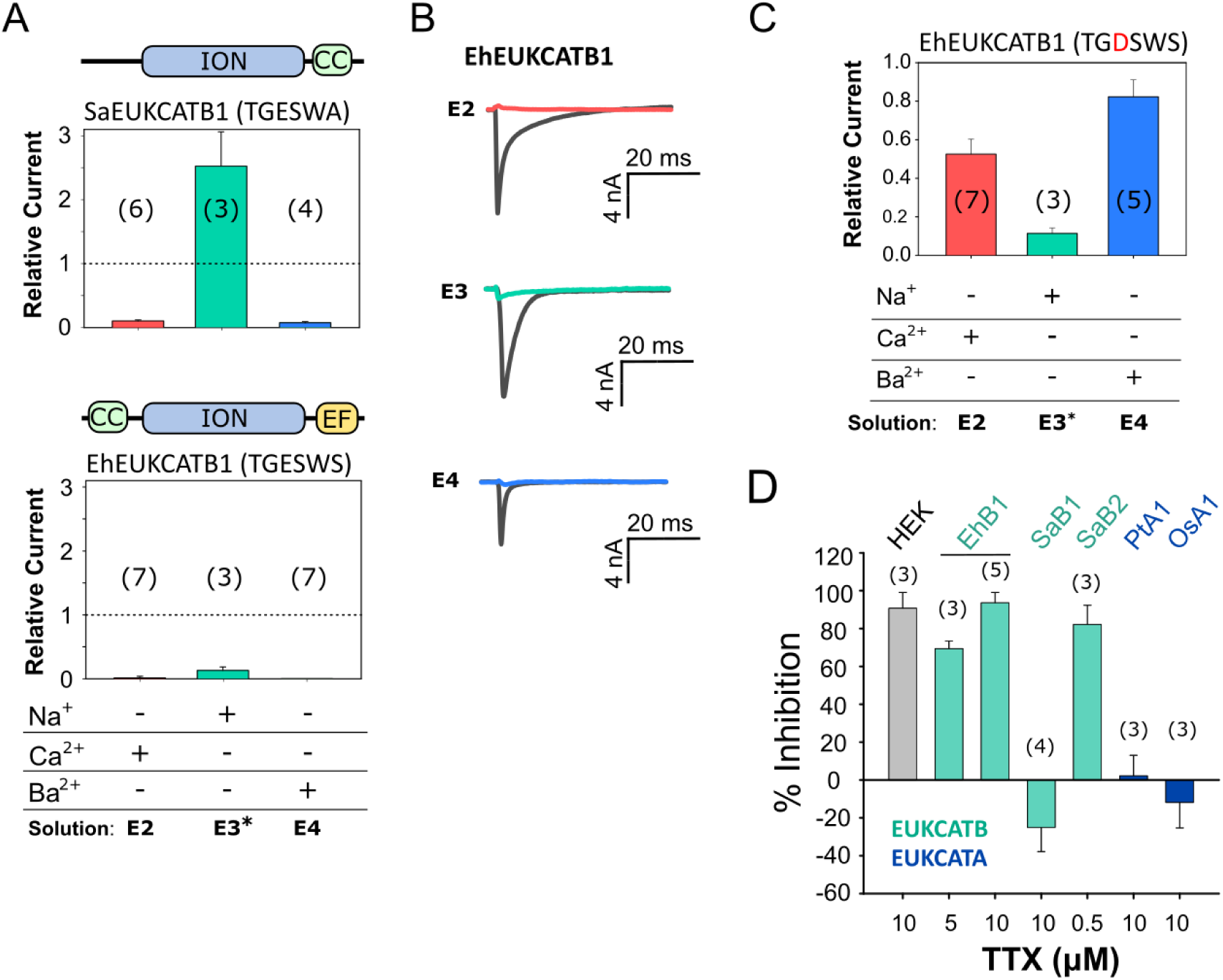
Coccolithophore EukCatB channels are Na+-selective. **A**. Mean peak currents for coccolithophore EukCatB channels (SaEUKCATB1 and EhEUKCATB1) are shown following substitution of different cations in the external solution. Currents are shown relative to the standard control external solution (E1; **Supplementary Table 4**). Replacement of extracellular Na^+^ with NMDG abolished the current in both EukCatB channels. Removal of Ca^2+^ did not reduce currents for SaEUKCATB1, but greatly reduced those of EhEUKCATB1, which contains two Ca^2+^-binding EF-hand domains at the C-terminus (*100 µM EGTA was added to EhEUKCATB1 experiments to ensure consistent removal of Ca^2+^). Schematic representation of domain architecture of each channel is shown above each graph; CC, coiled-coil domain (Arrigoni et al., 2016), ION, ion transport domain; EF, EF-hand pair. Error bars: SEM, *n* shown in parentheses. Extracellular and intracellular (pipette) solutions are given in Supplementary Table S4. **B**. Representative currents from (**A**) are shown for EhEUKCATB1. **C**. Mean peak currents following site-directed mutagenesis of the SF of EhEUKCATB1 from TGESWSE to TGDSWSE (E305D). Cation replacement experiments indicate that the EhEUKCATB1 E305D mutant conducts substantial currents in the absence of Na^+^ (i.e. when Na^+^ is replaced by NMDG in the external medium). The E305D mutation therefore causes the Na^+^ selective EhEUKCATB1 channel to become permeable to Ca^2+^ and Ba^2+^. Extracellular and intracellular (pipette) solutions are given in Supplementary Table 4, (*100 µM EGTA). Mean currents relative to control are shown (Supplementary Table 4). Error bars: SEM. **D**. The effect of TTX, a selective Na_v_ blocker, on the mean peak currents of EukCatB channels (*Emiliania huxleyi*, EhB1; *Scyphosphaera aptsteinii*, SaB1 and SaB2), compared to two EukCatA channels of diatoms (*Phaeodactylum tricornutum*, PtA1; *Odontella sinensis*, OsA1 (Helliwell et al., 2019). The effect of TTX on small native Na^+^ currents in HEK cells is also shown. Mean % inhibition (relative to untreated control current) is shown, error bars: SEM; *n* is given in parentheses.

The currents generated by EhEUKCATB1 were also dependent on the presence of external Na^+^. However, EhEUKCATB1 currents were absent in Ca^2+^-free media, suggesting that EhEUKCATB1 may represent a Ca^2+^-dependent Na^+^ channel (**Figure 3A and B**). EhEUKCATB1 possesses a C-terminal Ca^2+^-binding EF-hand pair that may contribute to the observed Ca^2+^-dependent activation. To confirm that the activity of EhEUKCATB1 is Na^+^-dependent, we used site-directed mutagenesis to modify its SF, changing the conserved Glu to Asp (E305D). The EhEUKCATB1-E305D channel was permeable to both Ca^2+^ and Ba^2+^ in the absence of Na^+^, indicating that it was no longer Na^+^-selective (**Figure 3C**). This result indicates that the highly conserved Glu (0) in the SF of coccolithophore EukCatBs is a critical determinant of Na^+^-selectivity in a manner resembling BacNa_v_ channels (DeCaen et al., 2014).

### The Na+-selective channel EhEUKCATB1 is TTX sensitive

Tetrodotoxin (TTX) is a highly selective and potent blocker of certain 4D-Na_v_1s (Ren et al., 2001; Klein et al., 2017), but single-domain BacNa_v_ are insensitive to TTX (Ren et al., 2001). We found that 5 µM and 10 µM TTX substantially inhibited EhEUKCATB1 **(Figure 3D**), lending strong support to this channel being a Ca^2+^-dependent Na^+^ channel. Although the sensitivity of EhEUKCATB1 to TTX is relatively low compared to the nanomolar sensitivity of some mammalian 4D-Na_v_s (Klein et al., 2017), it seems likely that specific structural properties shared by these channels are targeted by TTX. A direct comparison of the residues responsible for binding TTX between 4D-Na_v_ (pseudo-heterotetrameric) and single-domain EukCatBs (homotetrameric) awaits detailed structural information. Nevertheless, EukCatB channels possess a conserved Glu (E309 in EhEUKCATB1) (black arrow, **Supplementary Figure 3**), which is absent in TTX-insensitive BacNa_v_ and occupies a similar position to the ‘outer ring’ of charged residues in the pore of 4D-Na_v_ that have been implicated in TTX binding (Moczydlowski, 2013; Shen et al., 2018). However, we also examined the TTX sensitivity SaEUKCATB1. Unlike EhEUKCATB1, SaEUKCATB1 was insensitive to TTX, despite being highly Na^+^-selective **(Figure 3D**). By comparison, another highly similar EukCatB channel from *S. apsteinii* (SaEUKCATB2) was sensitive to 0.5 µM TTX. Thus as EukCatBs, like 4D-Na_v_s (Klein et al., 2017), are clearly not universally sensitive to TTX, further studies will be required to identify the residues that contribute to the different TTX-sensitivities in EukCatB channels. It should be noted that both of the two previously characterised Ca^+2^ and Na^+^ permeable EukCatA channels from diatoms (Helliwell et al., 2019) were insensitive to TTX.

### Coccolithophores exhibit fast electrical excitability

Given the presence of fast-activating Na^+^-selective voltage-gated channels in coccolithophores, we examined the nature of membrane excitability in these cells. Application of patch clamp approaches to decalcified *S. apsteinii* cells revealed fast-activating and fast-inactivating inward currents in response to membrane depolarisation (**Figure 4**). These extremely rapid kinetics are similar to those that underlie metazoan Na^+^-based action potentials, and indicate that *S. apsteinii* is an excitable cell capable of rapid electrical signalling. The currents also exhibit similar kinetics to those produced by the heterologously expressed SaEUKCATB1 channel (**Figure S2**). However, both *E. huxleyi* and *S. apsteinii* also possess 4D-Ca_v_/Na_v_ genes in addition to EukCatBs (**Table S1**) and so it is not clear which of these ion channels contribute to the observed membrane excitability. The coccolithophore 4D-Ca_v_/Na_v_ either exhibit the ‘EEEE’ motif that is associated with Ca^2+^ selectivity in 4D-Ca_v_1s, or a slight variant ‘DEED’ (**Figure 4D**) (Verret et al., 2010). As the presence of a lysine in the SF appears to be important for Na^+^-selectivity e.g. the DEKA of vertebrate 4D-Na_v_1s (Pozdnyakov et al., 2018), the coccolithophore 4D-Ca_v_/Na_v_ channels may not be Na^+^-selective, in contrast to the EukCatBs found in this lineage. Co-occurrence of both EukCatBs and 4D-Ca_v_/Na_v_s in coccolithophores may thus have driven functional diversification of voltage-gated ion channels. Investigation of the wider distribution and functional roles of EukCatBs and 4D-Na_v_/Ca_v_ in coccolithophores will provide insight into the selective pressures shaping the evolution of these different classes of ion channels.

**Figure 4.**
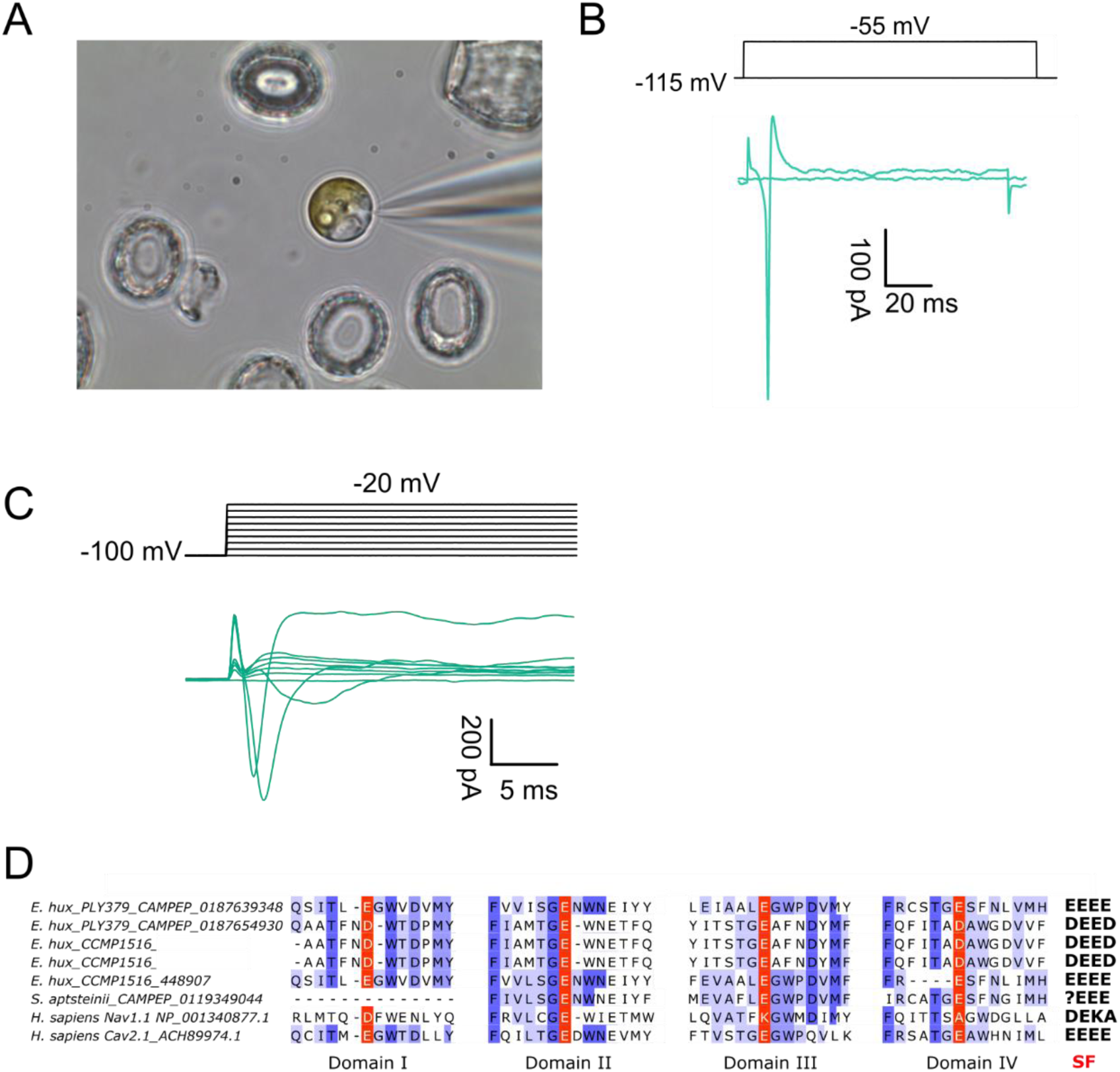
Fast voltage-activated inward currents in the coccolithophore *Scyphosphaera apsteinii*. **A**. Patch-clamp pipette forming a seal on a decalcified *S. apsteinii* cell (8.5 µm in diameter). **B**. Fast activating inward current evoked by a voltage command from a holding potential of −115 mV to −55 mV. Cell capacitance was 19 pF and seal was 5 GΩ. **C**. Current traces recorded in a decalcified *S. apsteinii* in response to membrane depolarisation from a holding potential of −100 mV in steps of + 10 mV. **D**. Multiple sequence alignment of the four pore domains of coccolithophore 4D-Ca_v_/Na_v_s. Sequences from coccolithophore genomes (*E. huxleyi* CCMP 1516) or transcriptomes (*E. huxleyi* PLY379 and *S. apsteinii*) are shown aligned to Na^+^ selective Na_v_1.1, and Ca^2+^ permeable Ca_v_2.1 from humans. The coccolithophore sequences most closely resemble the EEEE motif found in metazoan Ca^2+^ channels and lack conserved lysines found in Na^+^ channels. The 4D-Ca_v_/Na_v_ sequence found within the *S. apsteinii* transcriptome represents a partial sequence as it lacks domain I.

## Discussion

Coccolithophore EukCatBs represent a novel class of highly Na^+^-selective single-domain voltage-gated channels in eukaryotes. Their selectivity for Na^+^, together with their very rapid activation and inactivation kinetics, renders them distinct from the slower non-selective EukCatA channels recently identified in marine diatoms (Helliwell et al., 2019). Notably, many of the characteristics of the single-domain EukCatBs are highly similar to mammalian 4D-Na_v_s, particularly in terms of their activation and inactivation kinetics. Whilst the organisation of the four pore-forming domains into a single protein has likely enabled development of specialised structures and functions within 4D-Na_v_s (such as the intracellular linker connecting domains III and IV responsible for fast inactivation) (West et al., 1992), our results indicate that concatenation is not a pre-requisite for rapid gating kinetics, supporting observations from BacNa_v_ mutants (Irie et al., 2010; Shaya et al., 2014). Notably, certain coccolithophore EukCatBs also exhibited sensitivity to TTX, which is distinct from other single-domain channels such as BacNa_v_ and EukCatAs. However, as with mammalian 4D-Na_v_1s that are functionally divided into TTX-sensitive and TTX-resistant isoforms (Klein et al., 2017), we detected differences in sensitivity to TTX between EukCatBs. The EukCatBs may therefore prove to be useful models for studying Na^+^-channel gating and ion channel function more generally, as they are amenable to heterologous characterisation and exhibit key functional differences from BacNa_v_. The evolution of differing sensitivities of *S. apsteinii* EukCatBs also raises important ecological considerations. TTX is produced by a range of marine bacteria, and is also associated with harmful bloom forming dinoflagellates (Turner et al., 2017). Environmental exposure of coccolithophores to TTX could therefore have detrimental effects on EukCatB channel function, and consequently drive the evolution of resistant isoforms.

Na^+^ selectivity has arisen independently in animal Na_v_s at least twice (Gur Barzilai et al., 2012) and is thought to have been a critical step in the evolution of animal nervous systems, facilitating the rise of complex, fast moving multi-cellular organisms. The discovery of EukCatBs indicates that eukaryotes have also evolved alternative mechanisms to the 4D-Na_v_s for generating fast Na^+^-based action potentials. However, EukCatBs are not broadly distributed across eukaryotes and are unlikely to represent an ancestral channel in eukaryotes. Whilst the Na^+^-based signalling systems in animals are typically associated with the rapid transmission of information between distant cells of a multicellular organism, the requirement for fast Na^+^-based signalling in a unicellular algal cell is likely to be very different. Haptophytes do exhibit very rapid responses to stimuli, such as the coiling of their haptonema, a unique flagella-like organelle that aids capture of bacterial prey for phagocytosis by rapidly coiling or bending towards the cell body (Parke et al., 1955; Kawachi et al., 1991). Retraction of the 100 µm haptonema of *Chrysochromulina* sp. 21 NIES-4122 can occur within 5-10 ms (Nomura et al., 2019). However, given that the calcifying stages of coccolithophores from which action potentials have been recorded do not display a haptonema, broader roles are likely. Indeed, membrane excitability in coccolithophores may have a physiological role, as the major outward conductance at positive membrane potentials is H^+^, rather than K^+^ (Taylor et al., 2011). Repetitive action potentials in coccolithophores could therefore contribute to H^+^ efflux through H_v_1 channels and maintain pH homeostasis (Taylor et al., 2011).

The precise role of the EukCatBs in coccolithophores will require further investigation, although techniques for genetic manipulation are currently lacking in coccolithophores, hindering functional characterisation of individual genes in these organisms. Nevertheless, it is clear that the development of Na^+^-selective EukCatBs in haptophytes would have presented ancestral coccolithophores with the ability to uncouple electrical signalling from Ca^2+^ transport, as is the case in animal cells (Hille, 2001; Moran et al., 2015). Coccolithophores are unique amongst major calcifying organisms in that they produce their calcite structures (coccoliths) intracellularly, before they are secreted to the cell surface. Whilst intracellular calcification allows precise control of the environment for calcite precipitation, it also necessitates massive fluxes of Ca^2+^ across cellular membranes and highly specialised mechanisms for Ca^2+^ homeostasis. The presence of Na^+^-selective channels, uncoupling electrical signalling from Ca^2+^ influx, may have allowed coccolithophores to develop the unique mechanisms for Ca^2+^ homeostasis required for intracellular calcification, without disrupting cellular signalling.

In summary, we show that coccolithophores possess a novel class of ultra-fast voltage-gated single-domain Na^+^ channels, indicating that the capacity for rapid Na^+^-based electrical signalling is not unique to animals. Further characterisation of the structure and function of these unique Na^+^-channels is likely to inform us on the mechanisms underlying selectivity, gating and pharmacology of the wider voltage-gated ion channel superfamily.

## Materials and Methods

### Algal cultures

*Emiliania huxleyi* (CCMP1516) was obtained from the Plymouth Culture Collection (Marine Biological Association, Plymouth, UK). *Scyphosphaera apsteinii* (TW15) was from the Roscoff Culture Collection (Station Biologique de Roscoff, France). Cultures were maintained on a 12:12 light:dark cycle at 15°C with 100 and 50 µmol m^−2^ s^−1^ light. *S. apsteinii* was grown on medium that contained 90% filter sterilized Gulf Stream seawater (FSW) supplemented with Guillard’s f/2 nutrients and vitamins (Guillard and Ryther, 1962) and 10% K media (Keller et al., 1987), whereas *E. huxleyi* was grown on natural seawater supplemented with Guillard’s f/2 nutrients and vitamins (Guillard and Ryther, 1962). Cultures were maintained in mid-to-late exponential growth by sub-culturing 1 mL into 40 or 250 mL of fresh media every 2–4 weeks.

### Cell lines

HEK293 cells (ATCC CRL-1573) were grown at 37°C in 5% CO_2_ and 95% O_2_ in a humidified incubator. High glucose DMEM–Dulbecco’s Modified Eagle Medium growth medium with Antibiotic Antimycotic (Gibco), and 10% FBS (Gibco) was used to culture cells, which were passaged every 3 to 4 days at 1:6 or 1:12 dilutions (cell/mm^2^).

### Surveying for the distribution of single and four-domain voltage-gated ion channels

Sequence similarity searches were carried out as previously outlined by Helliwell et al., (2019) to survey haptophyte, pelagophyte and crypotophyte genomes and transcriptomes for single and four-domain voltage-gated ion channels. Query sequences from *Bacillus halodurans* C-125 NaChBac (protein id: BAB05220.1) and BacNa_v_-like sequences from the *E. huxleyi* CCMP1516 genome (protein id: 96075) (Read et al., 2013), in addition to the 4D-Ca_v_/Na_v_ sequence of *E. huxlyei* (protein id: 468996) were used. Transcriptome databases surveyed were from the Marine Microbial Eukaryote Sequencing Project (MMETSP, https://www.imicrobe.us/#/projects/104) (Keeling et al., 2014), alongside the reassembled sequence datasets (Johnson et al., 2018). In addition, eukaryote genomes including those from *E. huxleyi* (Read et al., 2013), *Pavlovales* sp. CCMP2436, *Aureococcus anaphagefferens* clone 1984, *Pelagophyceae* sp. CCMP2097, *Guillardia theta* CCMP2712, and *Bigelowiella natans* CCMP2755 were obtained from Joint Genome Institute http://genome.jgi.doe.gov/.

Databases were searched using BLASTP and TBLASTN with an E-value cut-off score of 1E^-10^. Each hit was inspected manually for relevant protein domains using Interpro (Apweiler et al., 2000), looking specifically for voltage-sensing domain (IPR005821), ion transport domain (IPR027359), EF-hands (IPR011992). The presence of a minimum of three pore domains were used as a threshold for candidate 4D-Ca_v_/Na_v_s in order to distinguish them from other voltage-gated cation channels. IDs of all protein hits reported in this study are given in **Supplementary Table 1**.

### Phylogenetic Analyses

The phylogenetic analyses of EukCat, Catsper and BacNa_v_ sequences were performed using multiple sequence alignments generated with MUSCLE via the Molecular Evolutionary Genetics Analysis (MEGA7) software (Kumar et al., 2016). After manual refinement, GBLOCKS0.91B was employed to remove poorly aligned residues, using the least stringent parameters (Castresana, 2000), resulting in an alignment of 172 and 211 amino acid residues for **Figure 1** respectively. Maximum likelihood trees were generated using MEGA7 with 100 bootstraps. Model analysis was performed in MEGA7 to determine an appropriate substitution model (WAG+G+I). Bayesian posterior probabilities were additionally calculated using BEAST v1.8.4 (Drummond et al., 2012) running for 10000000 generations.

### Synthesis of heterologous expression plasmids for HEK293 cells

Amino acid sequences of proteins used for heterologous expression are described in **Supplementary Table 2**. Coding sequences for EhEUKCATB1, SaEUKCATB1 and SaEUKCATB2 were obtained from MMETSP transcriptomic datasets: *E. huxleyi* CCMP 379 (CAMPEP_0187645740; MMETSP0994-7), and *S. apsteinii* RCC1455 (SaEUKCATB1: CAMPEP_0119314838, MMETSP1333; SaEUKCATB2: CAMPEP_0119345692, MMETSP1333). To confirm these sequences we amplified the open reading frame (ORF) from cDNA made from liquid cultures of *S. apsteinii* and *E. huxleyi* 1516 (using the primers: Sapsteinii_F1: ATGGTCGTCGCATCCTCAACG, Saptsteinii_R1: GCACCTCTGCACTCTGACATCTG and Ehuxleyi_F1: ATGATCGCGGCGATACATAAC, Ehuxleyi_R1: TCACACACGCTGCGTCGT). cDNA was synthesised using SuperScript III reverse transcriptase from RNA extracted using ISOLATE II RNA Mini Kit (Bioline) following the manufacturer’s instructions. Codon-optimised versions of the transcripts were then synthesised (GenScript, Piscataway, NJ) for characterisation in human expression systems, and sub-cloned into pcDNA3.1-C-eGFP using *HindIII* and *BamHI*. A 6 bp Kozak sequence (GCCACC) was included upstream of the ATG, and the stop codon removed.

### Site-directed mutagenesis of EhEUKCATB1

Site directed mutagenesis of the selectivity filter region of EhEUKCATB1 was performed using a Q5 Site-Directed Mutagenesis Kit (New England BioLabs, Hitchin, UK). Primer sequences (SDM1_F-TGACCGGAGAcTCCTGGTCTG; SDM1_R-GCACCTGAAACAGTGTGTAC) were designed (using the NEBase Changer Tool: https://nebasechanger.neb.com/), to replace a single glutamic acid residue with aspartic acid. Positive clones were screened by restriction analysis and confirmed by DNA sequencing of the entire gene.

### Transfection of HEK293 cells

HEK293 cells were plated for transfection onto glass-bottom (35mm) petri-dishes coated with poly-L-lysine (ibidi GmbH, Germany) to help with cell adhesion. Transfections of HEK293 were performed with 4 μL Lipofectamine 2000 (Invitrogen) and 1-2.5 μg plasmid DNA per 35 mm^2^, each prepared separately with Opti-MEM (Gibco). The Lipofectamine and DNA were mixed and allowed to rest for 5 min before 200 μL of the mixture was added to each plate. After 12-30 h of incubation, cells were rinsed and maintained with fresh growth media and kept in the incubator at 37°C with 5% CO_2_/95% O_2_ until used for electrophysiological experiments. Cells exhibiting GFP fluorescence were subsequently selected for electrophysiological analysis.

### HEK293 whole cell patch-clamp electrophysiology

Electrophysiological recordings were carried out at room temperature with an Axopatch 200B or Multiclamp 700B amplifier (Molecular Devices, Sunnyvale, California) through a PC computer equipped with a Digidata 1332 analog-to-digital converter in conjunction with pClamp 9.2 or pClamp10.1 software (Molecular Devices, Sunnyvale, California). Patch electrodes were pulled from filamented borosilicate glass (1.5 mm OD, 0.86mm ID) using a P-97 puller (Sutter Instruments, Novato, CA, USA) to resistances of 2-5 MΩ. In some experiments (analysis of SaEUKCATA1), unpolished electrode tips were coated with beeswax to minimize pipette capacitance. Data presented are leak subtracted and adjusted for liquid junction potential. Whole cell capacitance and seal resistance (leak) were periodically monitored during experiments. These corrected voltages were used to plot IV curves and in all subsequent investigations. The amplitudes of the currents were measured from the baseline to the peak value and were normalized for cell capacitance as whole-cell current densities (pA/pF). Steady-state activation was studied by measuring the peak sodium conductance (*G*_Na_) during a 50 msec test pulse to various test potentials from −120 mV holding voltage. *G*_Na_ was calculated from the equation: *G*_Na_ =*I*_Na_/(*V* −*V*_rev_), where *I*_Na_ is the peak sodium current during the test depolarization (V), and *V*_rev_ is the sodium reversal potential. Data were normalized to maximum peak conductance (*G*_max_) and fit to a two-state Boltzmann distribution: *G*_Na_/*G*_max_ = (1 + exp[(*V* −*V*_0.5_)/*k*])^−1^, where *V*_0.5_ is the potential for half-maximal activation, *k* is the Boltzmann constant.

To study steady-state fast inactivation, cells were held at prepulse potentials ranging from −140 to +30 mV for 50 msec and then subjected to a −10 mV test pulse for 10 msec. Normalized peak currents were plotted versus prepulse potentials, and curves were fitted by the Boltzmann function: *I*/*I*_max_= (1 + exp[(*V* −*V*_0.5_)/*k*])^−1^. Statistical analyses were performed with Sigma Plot 11.0 (Systat Software, Inc., Chicago, IL). Data are shown as the mean ± SEM. (*n*, number of experiments).

### Native phytoplankton cell patch-clamp recording and analysis

*S. apsteinii* cultures were harvested in mid-exponential phase for decalcification using a 0 Ca^2+^ artificial seawater supplemented with 20 mM EGTA (Taylor and Brownlee, 2003; Taylor et al., 2011). After 20-30 min cells were gently triturated with a transfer pipette to remove residual calcite and organic scales before plating onto a clean coverslip dish in full Ca^2+^ ASW. Whole cell patch clamp recordings were obtained with a pipette solution comprising (in mM) K-Glutamate 200, MgCl_2_ 5, EGTA 5, and HEPES 100, pH 7.5. Osmolarity was balanced to between 1000 and 1200 mOsmol using sorbitol. The extracellular ASW bathing solution contained (in mM) NaCl 450, KCl 8, MgCl_2_ 30, MgSO_4_ 16, CaCl_2_ 10, NaHCO_3_ 2, and HEPES 20, pH 8.0. Data were acquired using an Axon Instruments Axopatch 200B amplifier controlled through a Digidata 1200 with Clampex acquisition software (Molecular Devices, Sunnyvale, CA). Data presented are leak subtracted and adjusted for liquid junction potential. Whole cell capacitance and seal resistance (leak) were periodically monitored during experiments. Pipette resistance was 8 MΩ and whole cell capacitance varied between 12 and 22 pF among cells.

## Acknowledgements

We acknowledge support from the ERC grant ERC-ADG-670390 (C.B.), NERC Independent Research Fellowship grant NE*/*R015449/2 (K. E. H) and NSF grants 0949744 and 1638838 (A.T.).

